# Hydrophobic residues advance the onset of simple coacervation in intrinsically disordered proteins at low densities: Insights from field theoretical simulations studies

**DOI:** 10.1101/2021.01.18.427066

**Authors:** Satwik Ramanjanappa, Sahithya S. Iyer, Anand Srivastava

## Abstract

Intrinsically disordered proteins (IDPs) have engendered a definitive change in the way we think about the classical “sequence-structure-function” dogma. Their conformational pliability and rich molecular recognition features endow them with the ability to bind to diverse partners and predispose them to an elaborate functional armory. And of late, with studies on IDP-based liquid-liquid phase separation (LLPS) leading to formation of functional subcellular coacervates - best described as “membrane-less organelles (MLOs)”, IDPs are also bringing about paradigmatic changes in the way we think about biomolecular assemblies and subcellular organization. Though it is well recognized that the phase behavior of a given IDP is tightly coupled to its amino-acid sequences, there are only a few theories to model polyampholyte coacervation for IDPs. Recently, Joan-Emma Shea and co-workers used field theoretical simulations (FTS) to elucidate the complete phase diagram for LLPS of IDPs by considering different permutations of 50-residues chain representing 25 Lysine and 25 Glutamic acid [1]. Our work is an extension of that FTS framework where we develop and solve an augmented Hamiltonian that also accounts for hydrophobic interactions in the chain. We show that incorporation of hydrophobic interactions result in an advanced onset of coacervation at low densities. The patterning of hydrophobic, positive and negative residues plays important role in determining relative differences in the onset of phase separation. Though still very coarse-grained, once additional chemical specificities are incorporated, these high throughput analytical theory methods can be used as a starting point for designing sequences that drive LLPS.

## 1 Introduction

Intrinsically Disordered Proteins (IDPs) are functionally important despite lacking a native 3-dimensional fold [2–4]. Since they populate a range of conformations, they are best described by an ensemble rather than a single fold [5–8]. IDPs conformational landscape primarily deviates from a classical formula of sequence-based structure and function relationship generally applicable to folded proteins [2]. The conformational flexibility and multivalency of IDPs allows them to bind with diverse partners [9, 10] and make them multifunctional in nature [11, 12]. In living cells, these IDPs/IDRs-containing proteins undergo spatio-temporal separation leading to the formation of membrane-less organelles (MLOs) through multivalent interactions with other IDPs/IDRs, folded proteins, and nucleic acids [13–16]. In general, formation of these functional and dynamic biomolecular condensates require multiple proteins and nucleic acids molecules. However, there are studies where one IDP species is enough to form liquid droplets.

One of the fundamental questions in the field is how the phase-separation behaviour is encoded in the IDP/IDR sequences [17–21]. The relationship between amino-acid sequence distribution and ability of IDPs to form LLPS droplets have been explored extensively in the recent past using experimental, computational and theoretical approaches [22–25]. While the underlying design principles are still a matter of intense exploration, the one that stands out is the abundant presence of charged amino-acids in the sequence. Strategically placed positive and negative residues in these high charge content sequences generally account for high conformational entropy of the IDPs. And the shorter range interactions from the hydrophobic and aromatic residues that are interspersed across the sequence play important role in the formation of large stable and dynamic complexes. The delicate interplay between the above two potentially competing contributions [19, 26] namely the chain conformational entropy and the multiple interchain enthalpic interactions give rise to multivalent interactions between the IDPs that sometimes lead to LLPS and formation of liquid droplets.

IDPs typically interact with binding partners through short sequence motifs; short linear motifs (SLiMs), molecular recognition features (MoRFs), and low-complexity domain (LCD) sequences. While the role of charge content and patterning is undisputed, hydropathy patterning also plays an important role in dictating the multivalent nature and phase separation propensities of IDP complexes. In this context, LLPS in Tau protein is an interesting subject of study, where the LLPS is thought to be an intermediate state – capable of redissolving back into soluble state or transitioning into irreversible aggregates or amyloid fibrils [27–31]. A recent study shows that the fibril-prone and the reversible LLPS depends on the nature of interactions driving Tau condensation [32]. The study shows that Tau amyloid aggregation is favoured if the LLPS is hydrophobically driven while the electrostatically driven LLPS is largely reversible and aggregation-independent. The sequence patterning, evolved over millions of years, needs to be understood in terms of continuum of intermolecular interactions that ranges from electrostatic effects to shorter-ranged polar and dipole interactions, PI-orbital based interactions (PI-PI and cation-PI) and the hydrophobic effects, which are largely overlooked so far. Hydrophobic effects are emergent in nature and they are particularly important while the LLPS is underway and their role is not just limited to short-ranged stabilization of the already formed condensates.

Though it is well recognized that the phase behaviour of a given IDP is tightly coupled to its amino acids sequences, there are only a few theories to model polyampholyte coacervation for IDPs. The available analytical theories are largely derived from the rich field of polymer physics and are predominantly based on homopolymers. Given the importance of chemical specificity of the sequence and also the importance of solvation, computer simulations with detailed atomic models with explicit solvent would be ideal to test and/or predict LLPS propensities of a given solution. However, the length and time scales are formidable and computationally intractable even for a simple model systems [33]. Coarse-grained (CG) models, with reduced representations of amino acids and solvents, provide a computationally tractable alternative. And some recent CG simulations studies have been providing deeper insights into the relationship between polypeptide sequence and its phase behaviour [34–37]. With rising computational powers, faster sampling algorithms and better AAMD and CGMD forcefield parameters [36, 38–42], molecular simulations have the promise to unravel the design principles of sequence patterning that leads to LLPS in biomolecular solutions. Complementary to molecular simulation methods, purely analytical theories such as random phase approximation (RPA) [19, 43] and field theoretical simulations (FTS) [1, 44–49] go beyond simple mean-field approaches such as Flory-Huggins and Overbeek-Voorn theories [50]. These theories can be used with as high throughput first-pass methods to make broad-level predictions of the phase behaviour of IDPs. This is because these models take into account the chain connectivity and local fluctuations in polymer density, which makes them “sequence sensitive” and are theoretically well-grounded and can be formally derived by considering Gaussian fluctuations in the fields.

In general, classical self consistent field theory (SCFT) [44–49], a method widely used in polymer Physics field, becomes unreliable when there is significant density fluctuations and systems are close to phase transition. This often happens when one introduces electrostatic interactions and higher chemical specificity (detailed molecular interactions) on chains with limited degrees of polymerization. These inaccuracies can be avoided by using a fully fluctuating FTS models that promotes the Hamiltonian to complex values [44, 45, 51]. Though this leads to the famous “sign problem”, which can be effectively treated by adopting the complex Langevin (CL) sampling method widely used in nuclear physics [52,53]. In a wide-ranging and rich body of work, Glenn Fredrickson’s group has used FTS with great effect to study a variety of problems in polymer physics [44–49]. Recently, together with Joan-Emma Shea’s group, they have applied the method to elucidate the complete phase diagram for LLPS of IDPs where the authors have considered different permutations of 50-residues chain (25 Lysine and 25 Glutamic acid) and their study is focused on the role of charge-charge interaction in driving self-coacervation. Our work is an extension of the framework laid out in this recent publication where we develop an augmented Hamiltonian to account for apolar residues in the chain. Borrowing from the seminal work in the field of SASA and hydrophobic solvation calculation [54–58], we include an additional term in the Hamiltonian that is a function of solvent accessible surface area (SASA) and surface tension. These inclusion invokes approximations for making the functional integrals and partition function tractable for FTS. The new Hamiltonian and extended partition function makes the systems under consideration more realistic in their sequence decoration. We consider a variety of sequence with different patterns of positively charged, negatively charged and neutral apolar residues. To uniquely annotate the sequence decorations, we use a modification of scoring prescription propsed by Das and Pappu [59] to account for neutral residues in the score. Our work clearly shows how consideration of apolar residues (and hydrophobic solvation energy) affects the phase diagram and in particular the onset of the LLPS.

This article is organised as follows. In section 2 we describe the methods of formulation of hydrophobic interactions with which we score patters on chains that have both charged and apolar sites along with the field theoretic Hamiltonian and associated equations. After that we present and describe the key insights that we arrive at using our methods and how this can be used and refined to better models in other fields of biophysics and molecular organisations. In the appendix section more details of solving methods and how Gibbs ensemble was performed are described. All codes related to the methods development and example files are made available on Github (https://github.com/codesrivastavalab/fts-idpllps).

## 2 Methods

### 2.1 Sequence patterning with neutral beads and systems under consideration

We assign a unique score (*m*) to each of the sequence. The value of *m* helps in distinguishing the different patterns of positive, negative and neutral residues on the sequences. The scoring method used is an extension to “patterning parameter” κ introduced by Das and Pappu [59]. We modify the charge asymmetry parameter (*σ*) to include neutral residues. *σ* is now defined as 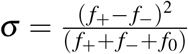, where *f*_+_, *f*_−_ and *f*_0_ are the fraction of positive, negative and neutral residues within the blob respectively. We find that a blob size of three instead of the commonly used size of five ensures that the uniqueness of the score in the sequences. The total number of blobs (*N*_*blob*_) is calculated by the number of sets of residues that is obtained by moving the window of size three with overlapping blobs. Asymmetry parameter is calculated for each sets of three residues. *δ* is the standard deviation of *σ* given as 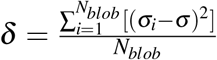 The sequence score *κ* is calculated by normalising 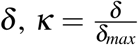. *m* ranges from 0 to 1 with κ = 1 for a system with a block polymer kind of arrangement of positive,neutral and negative residues.

For this study, we mainly consider two types of chains. In the first set of chains, we use only charge residues and the systems are same as the ones used by Joan-Emma Shea and co-workers [1]. We chose these systems both to validate our work as well as to show how the inclusion of explicit hydrophobic interactions in the FTS Hamiltonian affects the phase diagram even in case of pure polyampholytes. In the second set of chain patterning, we also consider neutral beads in the sequence. Here again, we have two broad class of sequence decoration. In the first set, we keep the neutral beads in the middle of the chain with flanking positive and negative beads and change the size of the central neutral band. In the second set, we have an interspersed neutral beads across the chain. These systems are designed to shed more light on the nature of phase diagram as a function of more realistic sequences as found in IDPs. Please see Fig. 1 for different patterning that we have used in this work.

**Figure 1:**
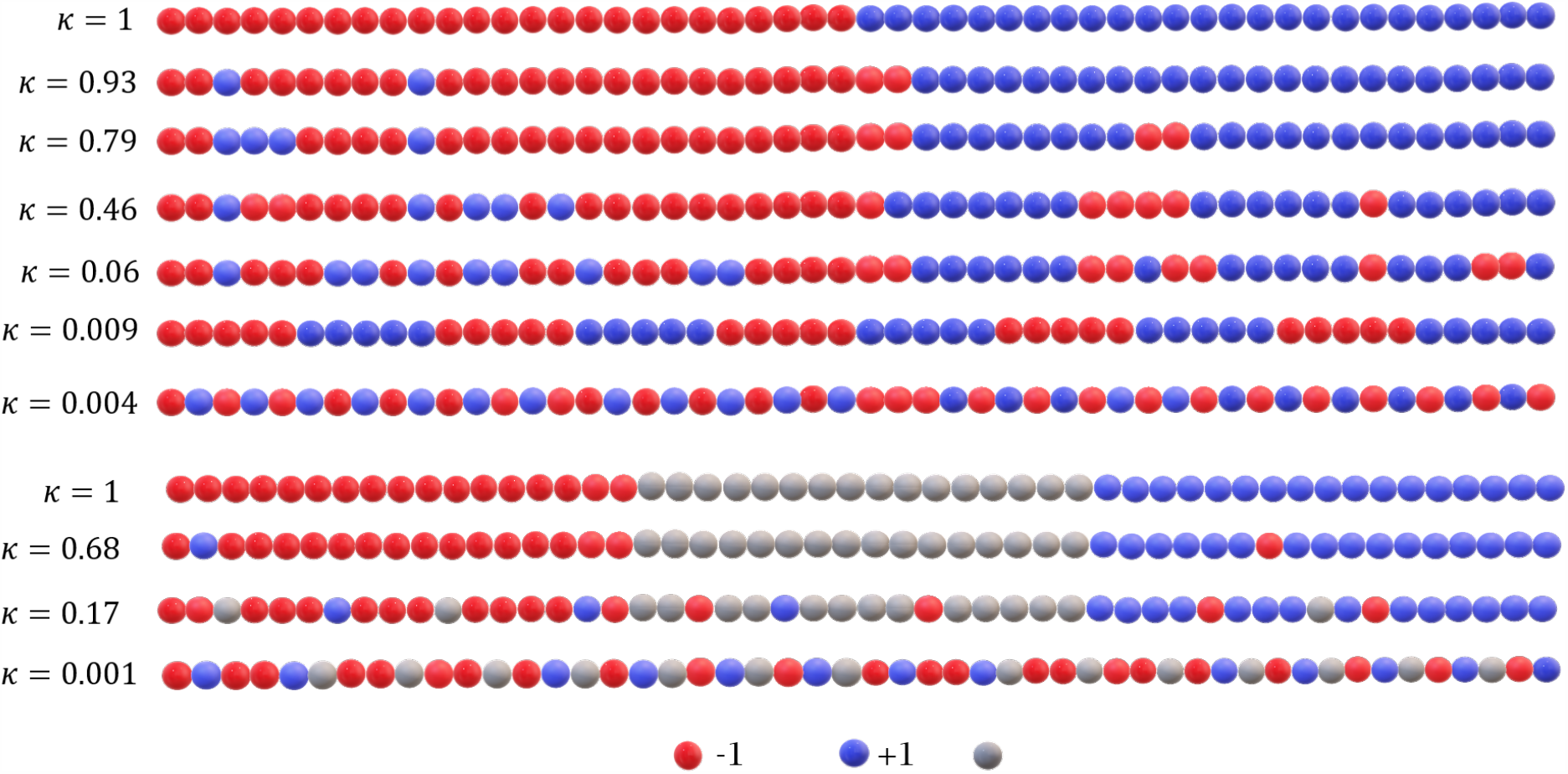
(Top panel) Value of m computed for various types of chains with no neutral sites as done in previous studies. (Bottom panel) Chain number for the chains with neutral sites as well, considered in following sections.

### 2.2 Theory and numerical implementation

#### 2.2.1 Hamiltonian for the fluctuating field theoretical model

For sake of completeness and to highlight the augmentation made to the existing model, we re-present an overview of polymer model with notations that are consistent with the original paper [1]. The system consists of *n* continuous Gaussian chains of volume *V* with *N* monomers per chain. This results in a monomer number density *ρ* of *nN/V*. Each monomer carries a monomer specific charge in terms of electronic charge *e* such that monomer *j* of chain *i* has a charge pattern *σ*_*i, j*_. The polymer chains are held together by a harmonic interaction potential given as:

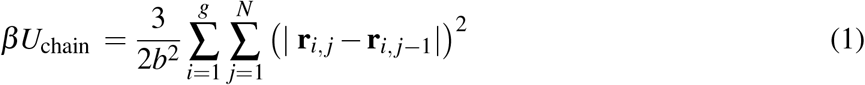

where *r*_*i, j*_ refers to the position of monomer *j* on chain *i, b* is the statistical segment length, and *β* =1*/k*_B_*T*. The monomers interact with a purely repulsive excluded volume interaction given as:

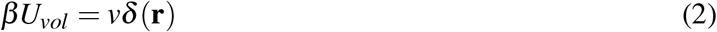

where *v* is the excluded volume parameter. Electrostatic interactions between charged monomers in a polymer chain as given as:

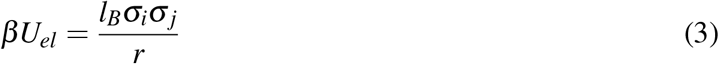

where *l*_B_ = *e*^2^*/* (4*πε*_0_*ε*_r_*k*_B_*T*) is the Bjerrum length, *ε*_0_ is the vacuum permittivity, *ε*_*r*_ is the relative dielectric strength of the solvent, and *e* is the fundamental charge of electron.

The total microscopic monomer number density and charge density are given as:

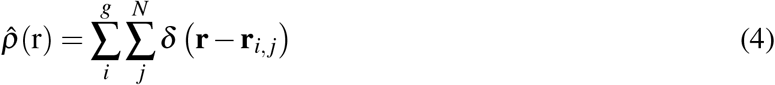

and

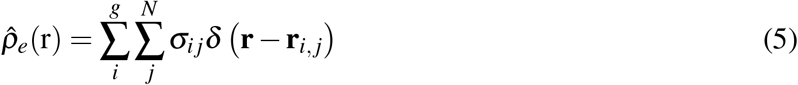

respectively. In order to maintain continuity, the number and charge density are smeared by performing a Gaussian convolution on Eqn 4 and 5. The smeared densities 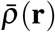 and 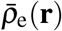 are given as:

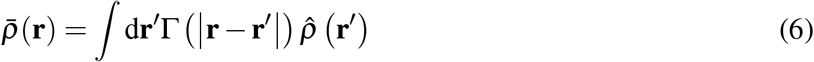

and

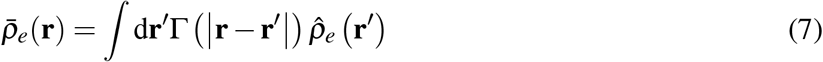

With

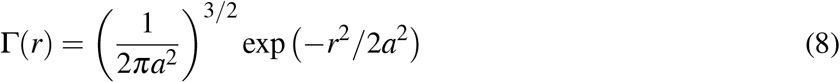

where “a” is normalised Gaussian smear width.

In this work, we add to the Hamiltonian an additional term to account for hydrophobic interaction. The formulation of the hydrophobic interaction terms come from approximations that make the formulations pliable for the FTS and we explain the same in greater detail in next section. The final form of the hydrophobic interaction potential that we arrive at is given as:

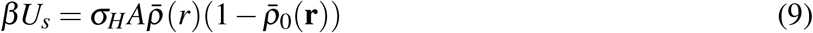

where *σ*_*H*_, scaled by *b*^2^, is the solvation surface tension term and A is the surface area of the monomer sphere. The value of *σ*_*H*_ determines the energy per unit surface area of the solute molecule exposed to a particular solvent. It is taken to be 0.6 J Å^−2^ (unscaled) for this work. In this work, we ignore the effect of hydrophobic interactions at very small distances. The Hamiltonian function has no particular form and the noise term also competes with this term leading to randomly staggered values.

Taken together, the total interaction energy Hamiltonian of the system written as the sum of bonded, excluded volume, Coulombic and hydrophobic interactions is given as:

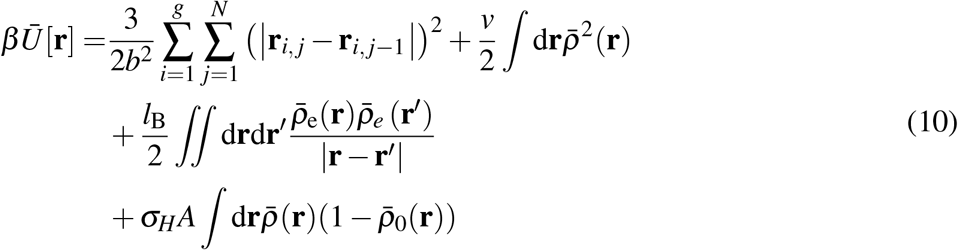

#### 2.2.2 Formulation details of Hydrophobic interactions

We use the the classical solvent accessible surface area (SASA) approach to account for hydrophobic interactions into the field theoretic hamiltonian framework. The choice of discrete Gaussian density makes it convenient to work with the integral form of the field theoretical approach. This formulation is mainly build using solid angle calculation and trigonometric manipulations and we demonstrate that in following text. For this work, we assume that all the molecules are spherical in shape. To illustrate the formulations, we will consider two different cases of solute-solute interactions. In the first case, two solute/monomer molecules are in close proximity. This is possible for two adjacent sites on a chain, two well-separated sites on the same chain making contact or intermolecular contacts. The maximum area of the solute molecule is blocked (due to presence of other molecule) from solvent when the molecule is in contact with the other spherical solute molecule. The magnitude of blocked area is computed by calculating solid angle area of angular separation the molecule creates on the considered solute molecule plus the angular separation the solvent molecule creates as shown in Fig. 2a. This would simply be given as 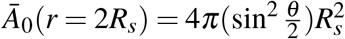 where *R*_*s*_ is radius of solute. Here, the area is expressed in terms of centre to centre distance between the solute molecules. For cases where there is a gap between proximal solutes that could allow a solvent space, we also require the information of blocked area for distances ranging from 2*R*_*s*_ to 2(*R*_*s*_ + *r*_*s*_), with *r*_*s*_ being the radius of solvent molecule as shown in Fig. 2b. The blocked area calculated as a function of distance between solute molecules is in such a situation is given as 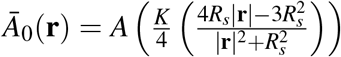, where *K* is sine squared of the angle subtended by solute and solvent molecule on the surface of considered solute as shown in Fig. 2b. This is expressed in terms of *R*_*s*_ and *r*_*s*_ where *A* is the surface area of a solute molecule equal to 4*πR*_*s*_^2^ and *Ā*_0_(*r*) is the blocked surface area due to single molecule of solute as a function of centre to centre distance between the molecules as shown in the Fig. 2. More details, in terms of the underlying algebra associated with the formulation, is provided in the following paragraph.

**Figure 2:**
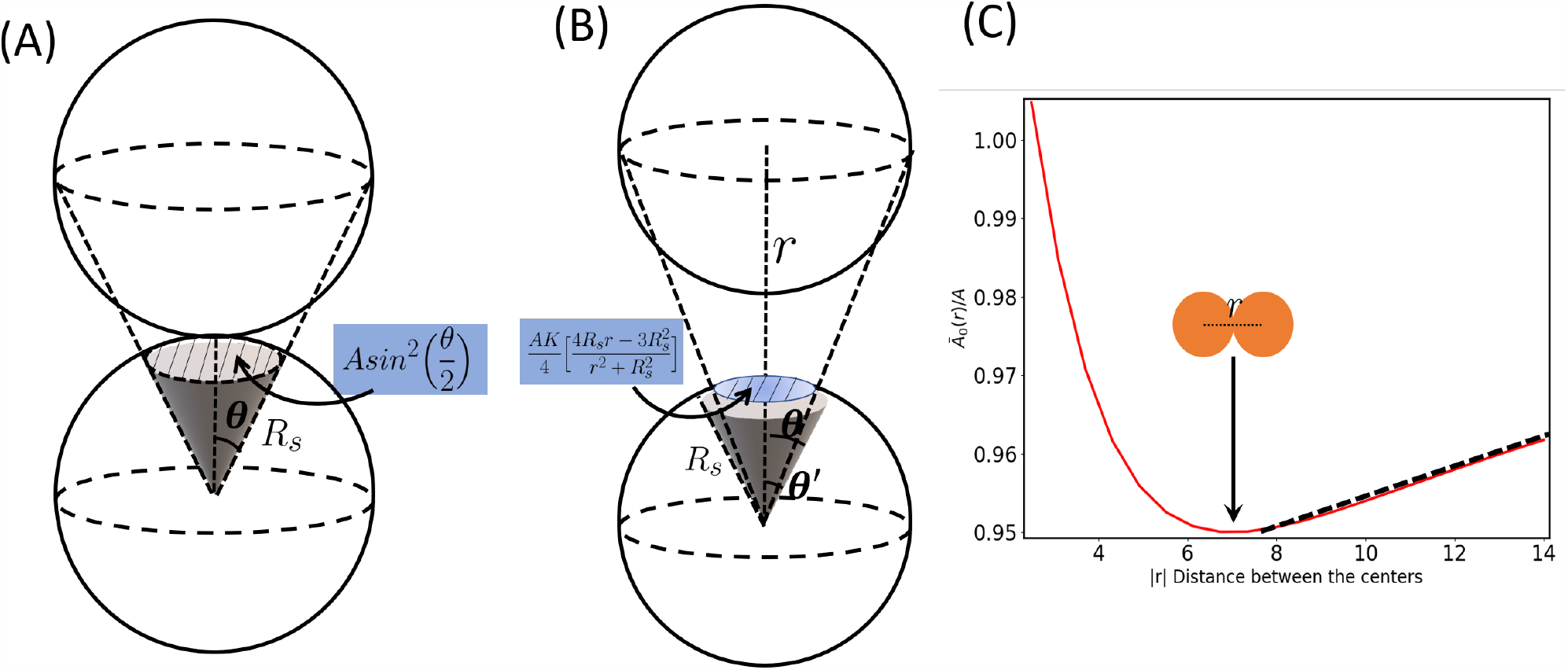
(Left panel) Case where maximum area is blocked by the solute molecule to interact with solvent. (Middle panel) Situation where magnitude of separation between the molecules is r. (Right panel) Function in Eqn 9 vs r by keeping 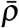 for simplicity of demonstration for *R*_*s*_ = 3.5Åand *r*_*s*_ = 2.5Å.

**Figure 3:**
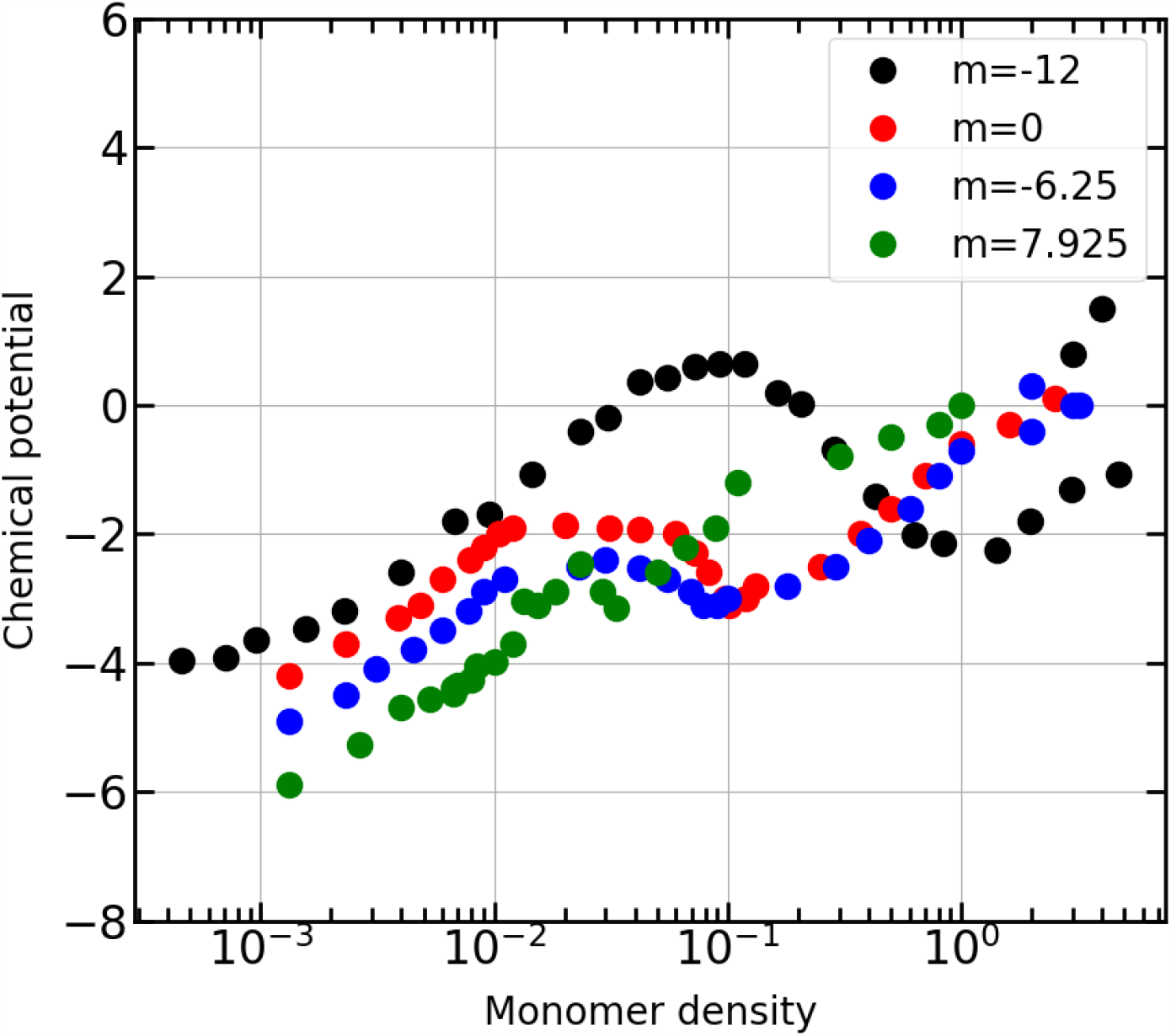
Chemical potential plots for various different charge decorated chains.

Starting from the simplest case for a solute placed over the surface of other as depicted in main text, the area blocked in this scenario is given by 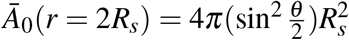. On calculating this sine of the angle in terms of *R*_*s*_ we get 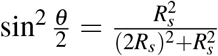. When we consider the second situation, where the proximal solutes can accommodate solvent molecules, we can take the difference in the blocked area from the first case to get the new blocked area as depicted in Fig. 2 and this is given as 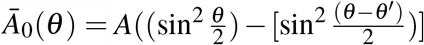. Here, we approximate 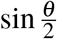 as 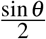 and also assume the limit of 90 degrees as the angle that is subtended by both the points at centre of the mobile sphere. On expanding the 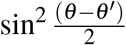 and putting back everything in terms of r and *R*_*s*_, for the sake of field theoretical formulation, we arrive at 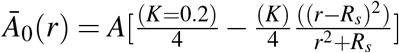, which can be rewritten as 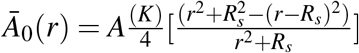. With this, we end up as the final form of 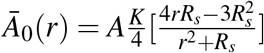.

The simple system of two solute molecules can then be used and generalised to normal system with inclusion of density of solute at that point to find the total contribution to blocked area due to those molecules and then add them over complete region. We find that the local fraction of blocked area can be computed using the density of solutes at that point as:

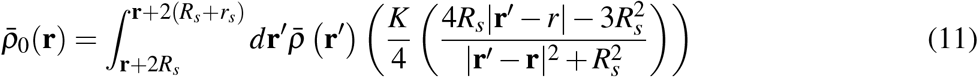

where 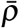 is the Gaussian smeared number density (Eqn 8). This form can be integrated over the whole region to get the total fraction of the blocked area blocked expressed as 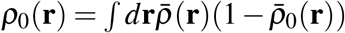 and the total surface exposed area for the whole system can be expressed as 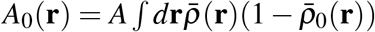 and incorporated in the full augmented Hamiltonian as in Eq. 10.

We subject the dependence of the blocked area to a particular range as the expression diverges for distance lower than the contact distance and becomes intractable in the integrable form. This approximation is acceptable since this distance has no influence on calculations of the blocked area. As shown above in equations above, formulation for computing SASA suggests complicated dependence on distance between molecules. This complication can be avoided for easier and convenient computational analysis by just considering the functional part only concerned with present work (i.e. the variation of this function for a range from 2*R*_*s*_ to 2(*R*_*s*_ + *r*_*s*_). The trend suggests that Fig. 2(c) (for *R*_*s*_ = 3.5Å) can be approximated to a linear function with definite slope and intercept. Values of slope and the intercept change depending on the value of (*R*_*s*_) and (*r*_*s*_), expression for *Ā*_0_ would become a simple dependence on r as 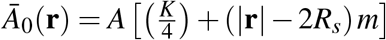. In the above demonstration value of *R*_*s*_ was taken to be 3.5Åfor which m was equal to 0.00168. For this work *R*_*s*_ was taken equal to 2.5Åfor which m was equal to 0.00123. The values of slope and intercepts to linearise can be found out by taking sufficiently good number of data points from original function and fitting it with a linear function. This kind of linear trend has reduced the the computational and theoretical expense to minimal. The simplified linearized expression for *Ā*_0_ is used in the entire work as an account to blocked surface area.

#### 2.2.3 The partition function and modified diffusion equation (MDE)

For our augmented Hamiltonian (Eq. 10), we write the partition function by keeping the ideal gas term as a constant and we introduce auxiliary field variables using functional integrals for excluded volume and electrostatic interactions terms as done by Shea and co-workers [1]. The auxiliary field variables for chemical potential field (*ω*) and electrostatic potential field (*ϕ*) are incorporated through Hubbard-Stratonovich transformation [60, 61]. We incorporate the hydrophobic interactions terms 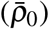 in the single chain formulation [62]. The partition function is expressed as:

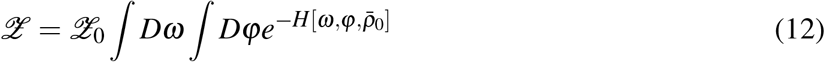

where 𝒵 is the partition function for an ideal gas with discrete Gaussian chains and other dependent parameters. The Hamiltonian, in terms of auxiliary field variables, is now rewritten as:

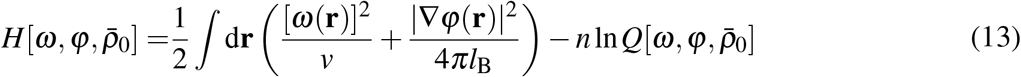

where 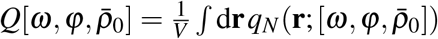 is a single chain partition function, which can be computed by the Gaussian chain propagator *q* as a solution to modified differential equation (MDE):

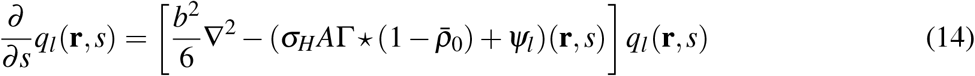

with initial condition *q*_*l*_(**r**, *s* = 0) = 1 and *ψ*_*l*_(**r**, *s*) entering Eqn 14 is given by *ψ*_*l*_(**r**, *s*) = *i*Γ *\** (*ω* + *σ*_*l*_*ϕ*), where 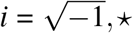 denotes a spatial convolution and *σ*_*l*_ is the charge of *l*^*th*^ monomer. 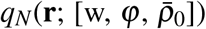 can also be constructed from Chapman-Kolmogorov equation [63]. The above differential equation was solved using operator splitting approach through discrete Fourier transform to get the initial configuration dependent field values. We also used the finite-difference approach and arrive at almost similar results for all cases. Field theoretical simulations are implemented by stochastically sampling the fields in complex Langevin dynamics where the variables *ω* and *φ* are propagated to complex space and given as:

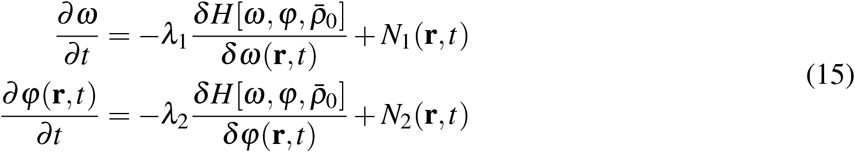

with

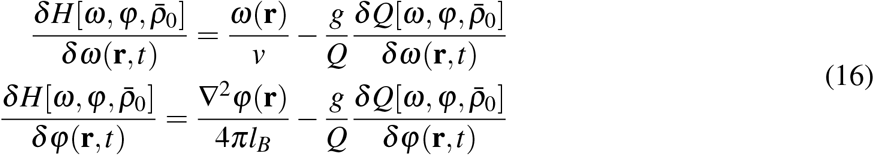

*N*(**r**, *t*) terms are real valued Gaussian White noise with zero mean and variance depending on dimensions of the simulation box, time stepping and volume segment. We update the field for every t = 0.02 time stepping in CL simulations. The modified diffusion equation (MDE) was solved with fixed contour stepping of Δ*s* = 0.01. We provide more details on numerical implementation for MDE and CL formulations in the next section. With partition function firmly in place, we proceed to calculated the various necessary quantities. Here chemical potential was computed with 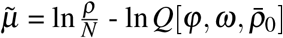 being chemical potential operator and *ρ* was the monomer number density. Numerical details of MDE and CL dynamics are discussed in Appendix. Also, following the CG dynamics calculations, we use the regular Gibb’s ensemble method to arrive at the phase coexistence curve. For sake of completeness, Gibb’s ensemble workflow is also presented in the Appendix.

## 3 Results and Discussion

In both molecular simulations and analytical theory work, role of charge patterning in IDP chains has been extensively studied to elucidate the conformational preferences and phase separation behaviour of the IDP. There has been relatively little progress in accounting for the role of other types of interactions such as hydrogen bonding interactions, weak non-covalent interactions (*π* − *π*, Cation-*π*)and hydrophobic interactions. In this work, we explore the effect of hydrophobic interactions and as well as hydrophobic residues patterning on the phase co-existence behavior.

### 3.1 Effect of hydrophobic interactions on phase coexistence

The effective charge distribution in sequences on the binodal curve was studied extensively by Shea and co-workers [1]. Sequences with different degree of “patchy” distribution of charges, quantified by the *κ* values showed an increase in critical Bjerrum length with decrease in *κ* of the sequences. A direct comparison to the experimental phase diagrams can be made by considering the inverse relation of Bjer-rum length and temperature. The sequences show increasing upper critical solution temperatures with increase in the *κ* values. This behaviour is driven by the predominance of intra-sequence electrostatic interactions in sequences with larger “patches” of contiguously charged residues. In Figure 4, we show the effect on the binodal when hydrophobic interactions is considered for a polyelectrolyte where we show the phase diagram for two sequences at extremes of the charge decoration “patchiness’. The points on the coexistence curves were determined using Gibb’s ensemble [46] method as described in the methods section. While the behaviour of increasing upper critical solution temperature of sequences with higher *κ* score is preserved with the inclusion of hydrophobic interactions, the onset of phase separation occurs at lower densities. This can be attributed to higher contribution of the hydrophobic interactions than the electrostatic interactions to the Hamiltonian at lower densities. Since field theoretical simulations do not provide information about geometries of the sequences, it is difficult to comment on how addition of hydrophobic interactions effect single chain collapse at lower densities.

**Figure 4:**
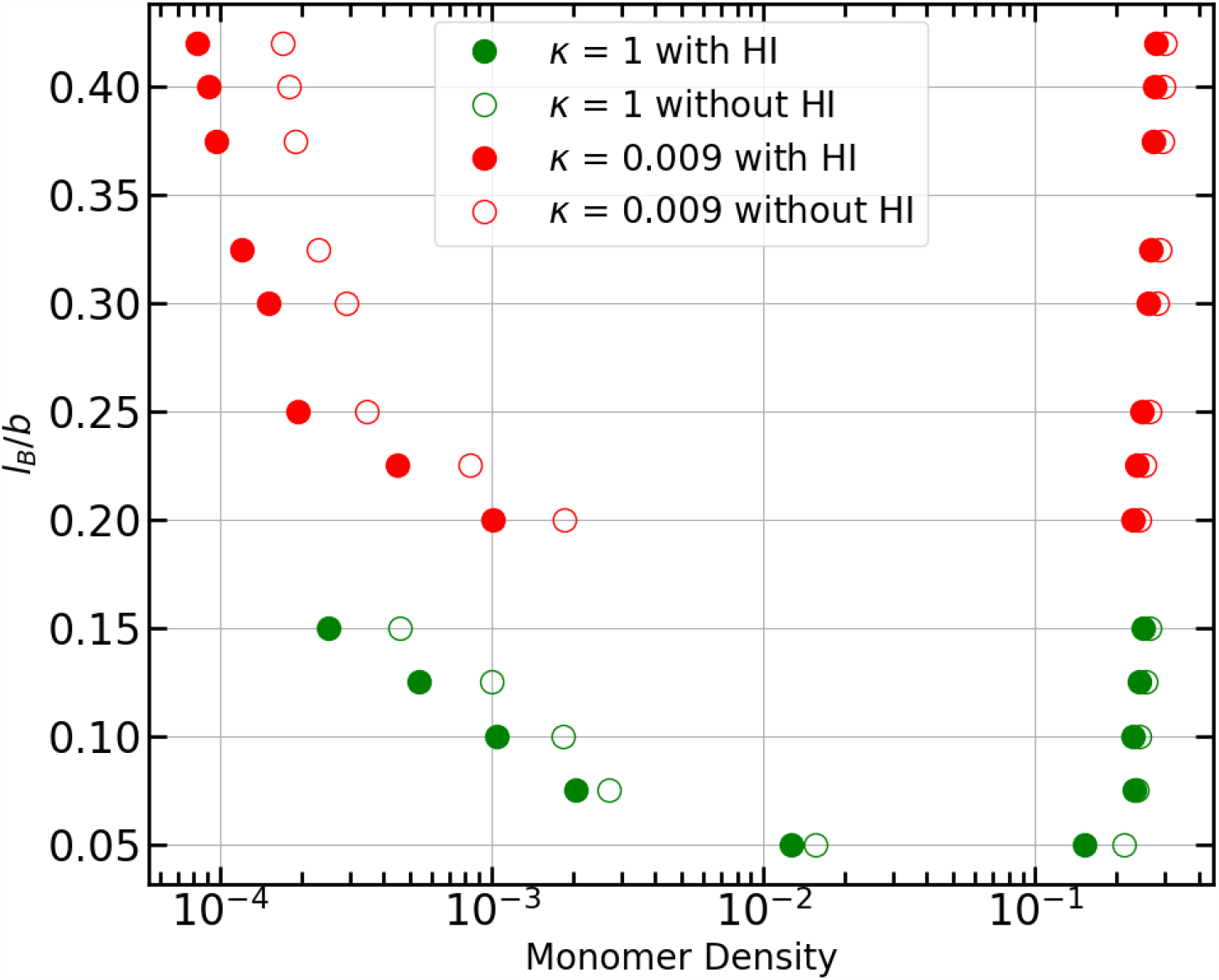
Phase diagram for sequence with lower and higher κ values. Lower κ have higher critical Bjerrum length. Sequences corresponding to different *κ* values is shown in Figure 1.

**Figure 5:**
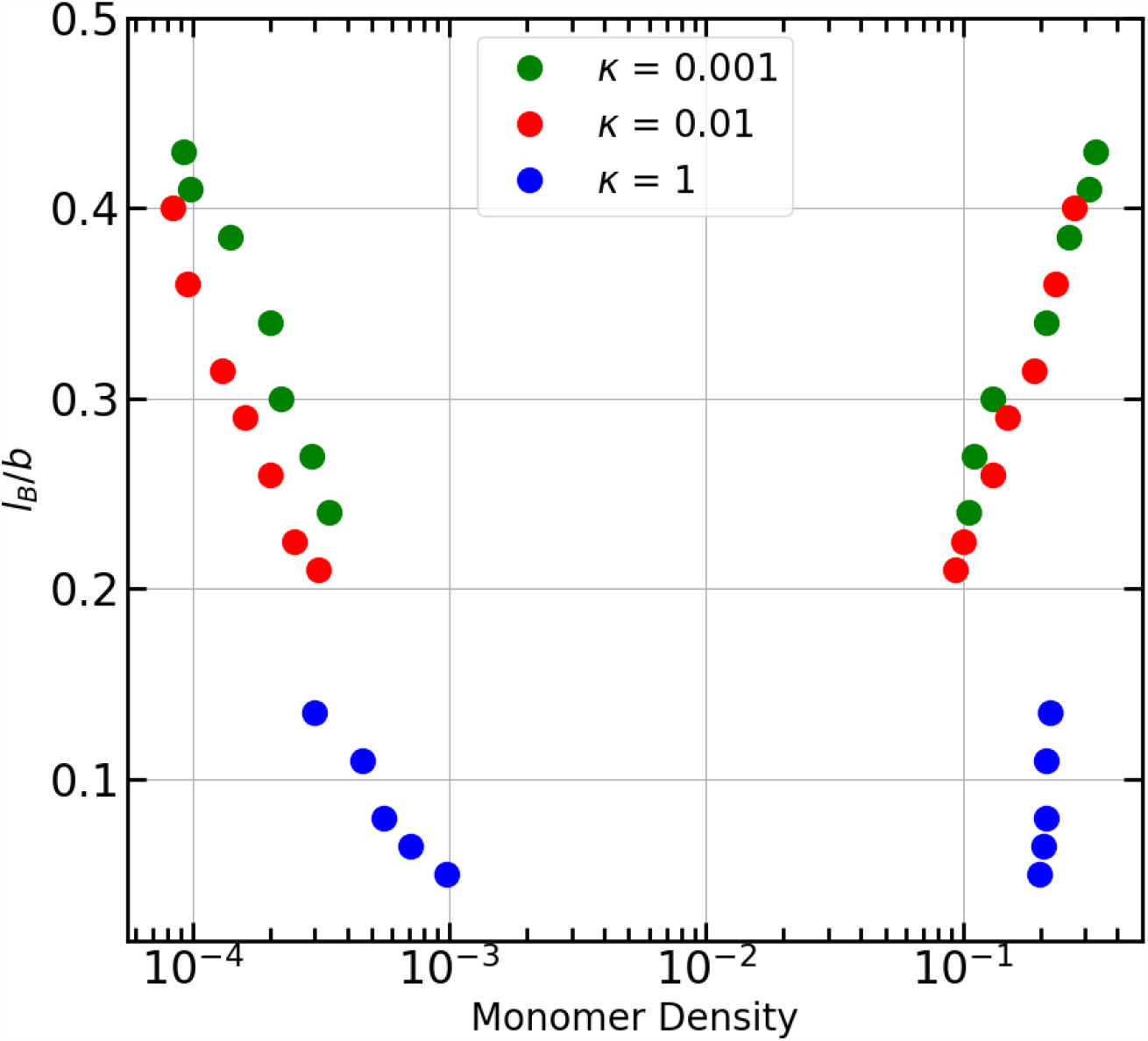
Phase diagram for sequence with lower and higher κ values. Lower κ have higher critical Bjerrum length. Effect of HI is seen more evident in systems enriched with contiguous neutral patches.

### 3.2 Effect of inclusion of hydrophobic interactions in binodal of sequences with apolar residues

The role of hydrophobic interactions in self-coacervation leading to LLPS has been studied extensively in elastin-like peptides and tau protein [64–66]. The effect of inclusion of hydrophobic interactions is more evident in sequences that have apolar sites. These sites do not have an electrostatic interaction term, hence making the hydrophobic interactions the only other non-bonded interaction term in the Hamiltonian governing sequence organization. In the following section, we explore the effect of different hydrophobic patterning as well as different hydrophobic stretch-size on the phase co-existence behaviour.

#### 3.2.1 Different decorations

Similar to the effect of “patches” of distribution of charge beads on the critical Bjerrum length, the decorations of apolar residues also determine the critical Bjerrum length. Figure 6 shows the phase diagram of sequences with different patterns of charged and neutral beads arrangement. The arrangement of sequences with κ value of 0.001, 0.1 and 1 are shown in Figure 1. All these sequences have conserved number of positive, neutral and negative beads. The critical Bjerrum length is higher for sequences with lower κ values. Bjerrum length can be an indicator of tempearture of the system to some extent though we note that a direct connection between Bjerrum length and temperature cannot be made in these systems due to the inclusion of hydrophobic interaction term in the Hamiltonian.

**Figure 6:**
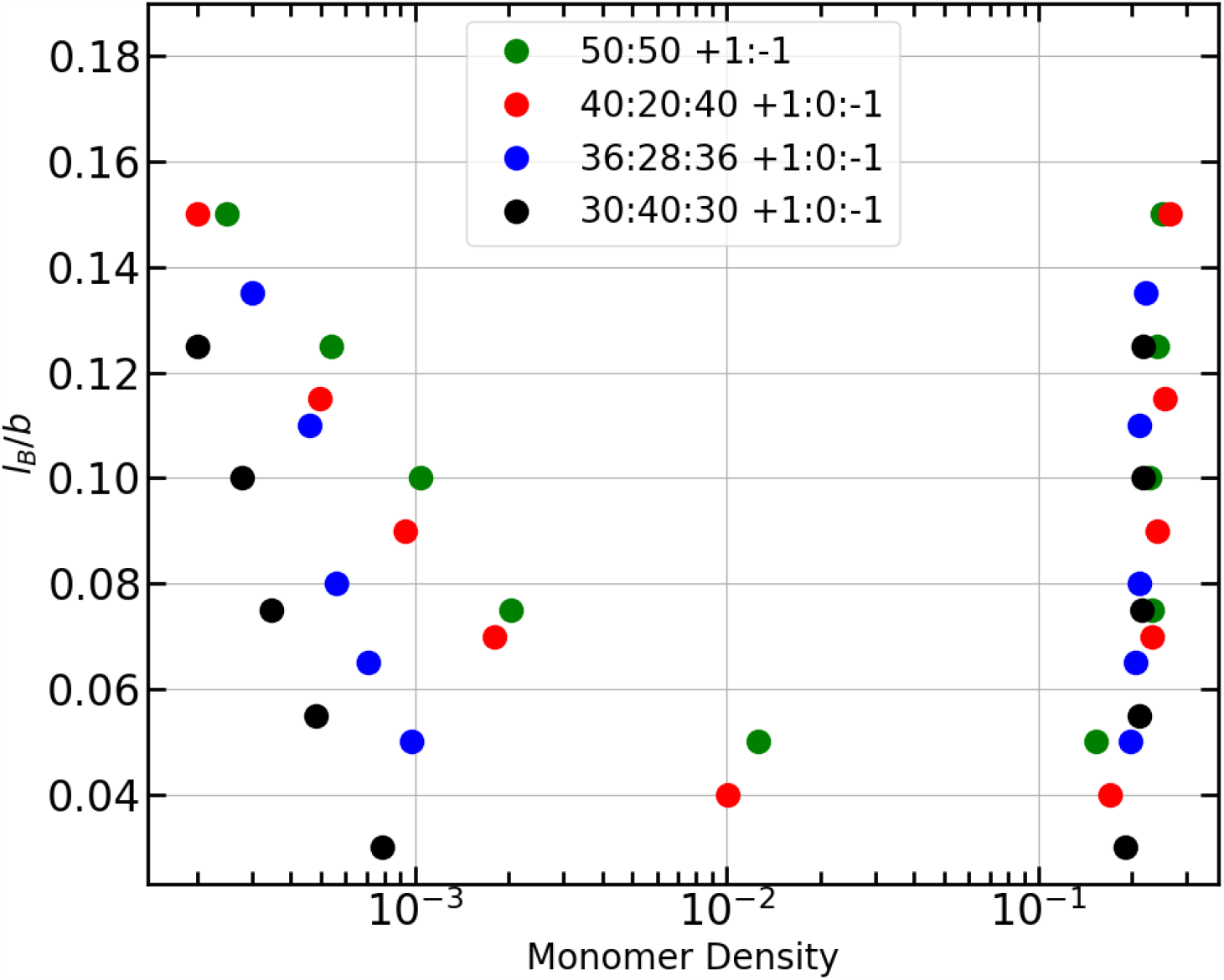
Phase diagram for triblock polymers with different lengths of stretches of neutral residues for sequences with same chain length

Apart from the effect on critical Bjerrum length, inclusion of hydrophobicresidues also result the onset of LLPS at lower monomer densities. This is similar to the effect of inclusion of hydrophobic interactions in sequences with no neutral residues. In Figure 6, the phase diagram obtained for sequences without hydrophobic interaction term is shown in open circles. The onset of LLPS at lower monomer densities for sequences after accounting for hydrophobic interactions is evident from the figure. Furthermore, the effect of hydrophobic interactions on the onset of LLPS at lower densities has more effect on polymer chains with less charge decorations i.e. block type polymers than in distributed systems.

#### 3.2.2 Different stretches of hydrophobic centers

We study another aspect of inclusion of neutral residues with hydrophobic interactions by varying the length of apolar neutral block in the triblock system while keeping the total chain length same. As expected, with increase in length of the neutral hydrophobic stretch, the onset of LLPS occurs at lower densities. The possible entropic origin of these self-coacervation at lower densities is better examined using particle based simulation studies. Though an extensive study of low-complexity regions like the RGG/RG motifs in driving LLPS has been documented [67, 68], to the best of our knowledge, the nature of length of hydrophobic amino acid motifs that drive LLPS is yet to be thoroughly studied.

Generally, a diblock polymer of opposite charges always tends to be in a hair pin like structure. On the contrary, polymers with distributed charges tend to be in a coiled state and are more collapsed. One would expect similar behavior in IDPs. Chains with distributed charges are in considerably collapsed or coiled state than the ones with stretches of like-residues in blocks. For a noticeable hydrophobic stretch, SASA contribution becomes more effective for phase separation to happen. This can also be seen from the changes in chemical potential with concentration as seen in Figure 6, which clearly portrays unique behavior of triblock chain, (black dots) with that trend fading with more distributed systems. As a result the changes observed in later system is less.

#### 3.2.3 Different values of solvation coefficient

Apart from Bjerrum length, solute-solvent interactions governed by exposed surface area, also determine the coexistence points on the binodal. As per the definition, with larger value of *σ*_*H*_, the chains would avoid exposure to the solvent resulting in faster phase separation. This was verified by increasing the value of *σ*_*H*_ in above calculations. Figure 7 shows that an increase in *σ*_*H*_ results in the shift of phase boundaries to to lower densities.

**Figure 7:**
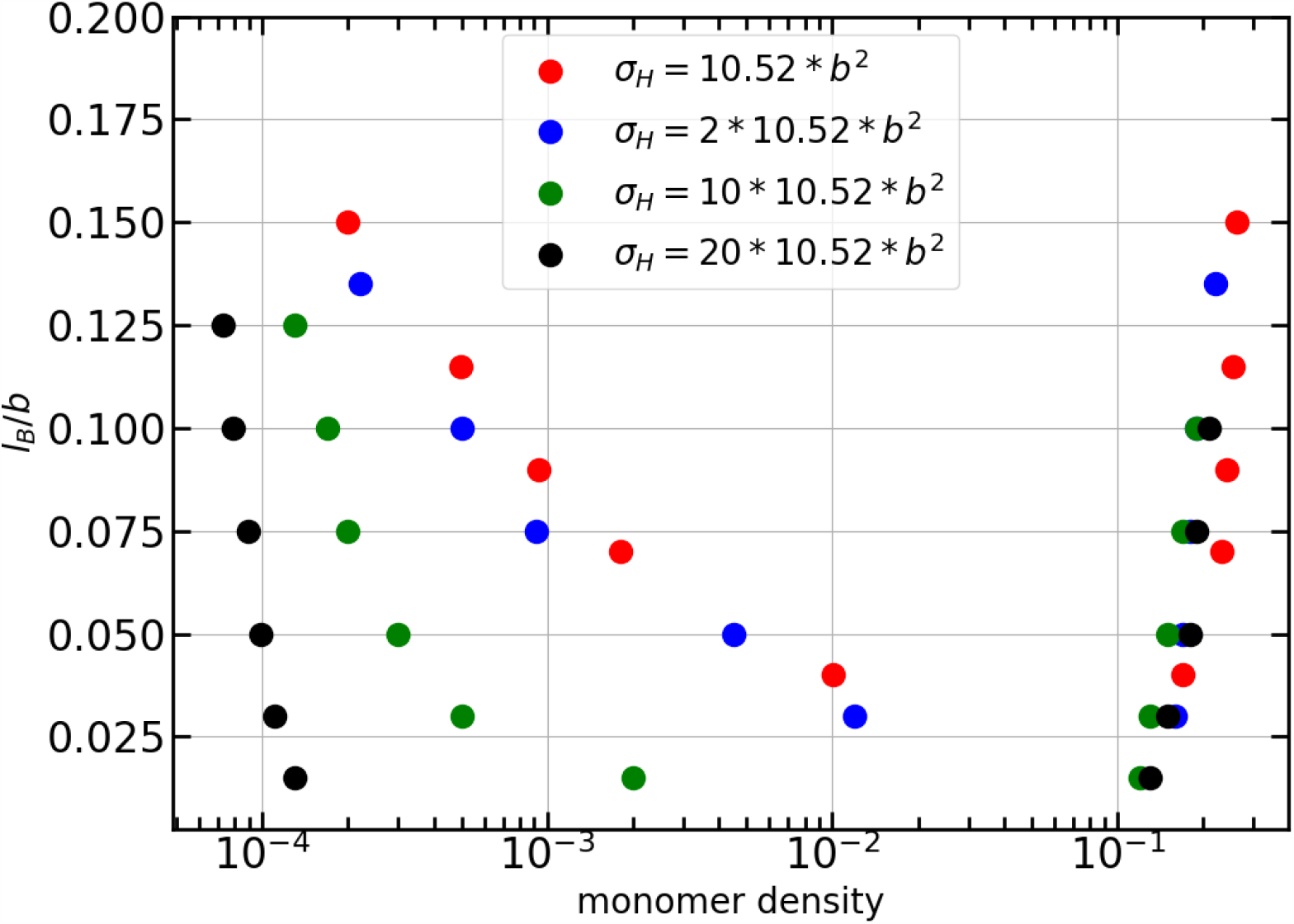
Phase diagram for triblock polymers with different *σ*_*H*_.

## 4 Conclusion and Outlook

In this work, we have attempted to describe phase coexistence with incorporating hydrophobic interactions through SASA approach, which gives us better understanding of LLPS for studies in biology. This entropically driven effect with more sophisticated studies can can give biologists a better understanding of formation of cell organelles and biological globules formed by IDP’s. We would need better formulations for hydrophobic interactions for more intrinsic details, as seen earlier we have ignored the contributions from lower distances which FTS for this formulation fail to give accurate values, better models need to address this issue.

## 5 Appendix

### Numerical details of MDE and CL dynamics

We first calculate the initial values for the auxiliary fields through Fourier transform by moving into the reciprocal space. We solve MDE using operator splitting method (as shown below) as well as well known finite difference algorithm which happen to agree within appreciable error. In case of operator splitting algorithm, the formal solution to the diffusion equation is expanded to second order in monomer step Δ*s* such that:

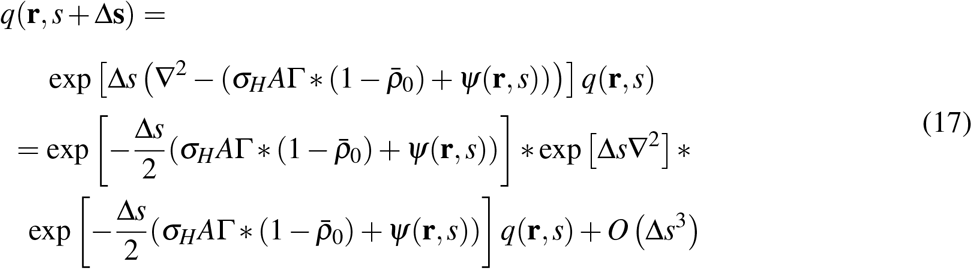

The solution of the diffusion equations indeed represents the most computationally demanding component of FTS. Runge-Kutta algorithm was used to propagate CL equations with time step of Δ*t* = 0.1. The chemical potential operator as described in the main text was used to compute chemical potential and pressure operator as by Shea and co-workers [1] with small modifications was

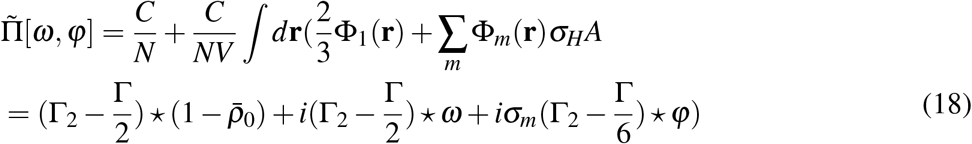

where C is the scaled monomer density (*ρb*^3^) and *\** denotes a spatial convolution. Γ_2_(**r**) = (1 − *r*^2^*/*3*a*^2^ Γ(**r**) with Γ(**r**) a normalized Gaussian. In the above equation Φ_1_(**r**) is given as 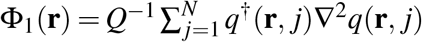 where *q*(**r**, *j*) is the forward Gaussian chain propagator, and *q*^†^(**r**, *j*) is a similar reverse chain propagator, and Φ_*m*_(**r**) represents the concentration of all the non interacting fields Φ_*m*_(**r**) = *Q*^−1^ ∑ _*j*∈*m*_ *q*^†^(**r**, *j*)*q*(**r**, *j*), which is a monomer-type *m* density field computed by counting monomer segments of monomer-type *m*. Here in this work, we used only three types of monomers *σ* = −1, 0, 1.

### Gibbs ensemble and phase separation equilibrium

After the CL runs, we constructed the binodal curves by performing FTS in Gibbs ensemble. This is conventionally done by partitioning the polymer in solution of total volume V into two phases of *V*_*I*_ and *V*_*II*_ with corresponding number of chains *g*_*I*_ and *g*_*II*_, respectively such that *V*_*I*_ + *V*_*II*_ = *V* and *g*_*I*_ + *g*_*II*_ = *g* [1]. We arrive at the equilibrium condition when the chemical potential and the osmotic pressure of both phases are equal. In Gibbs ensemble, the total simulation box of volume V is partitioned into two regions corresponding to a dense and dilute phase. The concentration and phase volumes change continuously till we reach equilibrium between the two phases. We scaled all the lengths by 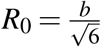 with b as segment length and reduced chain density 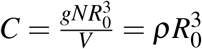, which we used to plot phase diagrams. From the equation of balances, we get *C*_*T*_ = *v*_*I*_*C*_*I*_ + *v*_*II*_*C*_*II*_ with *C*_*T*_, *C*_*I*_*andC*_*II*_ as total chain density, chain densities of region I and II, respectively. *v*_*I*_ and *v*_*II*_ are the volume fractions defined as *v*_*I*_ = *V*_*I*_*/V* and *v*_*II*_ = 1 − *v*_*I*_. Differentiating with time and with concentration rate proportional to difference in chemical potential of either medium and similarly correlating between the rate of change of volume and pressure change, we get:

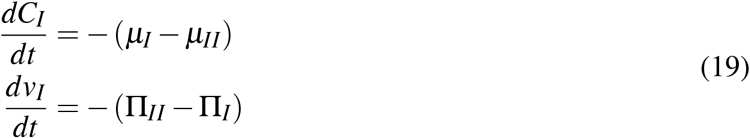

which can be solved numerically by simple Euler stepping algorithms. We used Runge-Kutta methods to solve and update the scheme as:

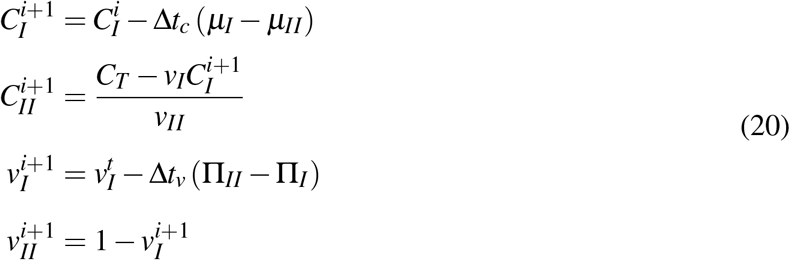

In our work, we maintain charge neutrality and we run our simulations for 10^6^ CL steps, updating concentrations and volume according to above equations for every 1000 steps with Δ*t*_*c*_ = 0.05 and Δ*t*_*v*_ = 2. We recorded the concentrations *C*_*I*_ and *C*_*II*_ at the completion of Gibbs ensemble runs as this dataset form the points of boundary for the binodal curve for particular temperature in the phase diagram.

## Acknowledgments

A.S. thanks the Indian Institute of Science and the Ministry of Human Resource Development of India for the start up grant and the Department of Science and Technology of India for the early career grant. This research was also supported by the Department of Biotechnology, Government of India in the form of IISc-DBT partnership programme. Support from FIST program sponsored by the Department of Science and Technology and UGC, India – Centre for Advanced Studies and Ministry of Human Resource Development, India is gratefully acknowledged by the authors.

## Notes

### Competing Interest Statement

The authors have declared no competing interest.

